# Temporal evolution and spatial heterogeneity of cerebral cortical-depth profiles of the BOLD-fMRI response

**DOI:** 10.1101/2025.09.18.677117

**Authors:** Anna I. Blazejewska, Daniel E.P. Gomez, Jonathan R. Polimeni

**Author notes:** Correspondence should be directed to: Anna Blazejewska, Ph.D., Athinoula A. Martinos Center for Biomedical Imaging, 149 13^th^ St. Suite 2301, Charlestown, MA, 02129 USA. Richard M. Lucas Center for Imaging, Department of Radiology, Stanford University School of Medicine, Stanford, CA, USA.

## Abstract

Functional MRI measures brain activity by tracking the associated hemodynamic response, which is shaped by the vascular anatomy. Microscopy studies have shown that the hemodynamic response initiates within the cerebral cortex then spreads upwards to the pial surface, where the largest fMRI signal changes are seen, however responses at the cortical surface exhibit poor neuronal specificity. Motived by this, we characterized the time-evolution of the fMRI response in humans. At some cortical locations the fMRI response peaked at the pial surface but in others it peaked within the cortex, likely due to the sparsity of large pial vessels and the dense capillary bed in middle cortical layers. We observed the earliest response onset in middle cortical depths, which is also the site of thalamocortical input. Standard approaches that aggregate fMRI responses over space and time may therefore lose meaningful information, and “physiologically-informed” strategies may enhance neuronal specificity of fMRI measurements.

## INTRODUCTION

Although functional Magnetic Resonance Imaging (fMRI) is the most widely used method for measuring brain function noninvasively across the human brain, all fMRI techniques in use today measure brain activity by tracking associated hemodynamic responses that promptly deliver fresh blood to active neurons. This is often viewed as the fundamental limitation of the technique when applied to measuring brain function (Logothetis, 2008). However, mounting evidence mainly from invasive in-vivo microscopy studies in small-animal models suggests that the precision of blood flow regulation in response to neuronal activity in the cerebral cortex is remarkably high, and blood flow is controlled at spatial and temporal scales smaller than what can be achieved even with modern MRI technologies (Polimeni and Wald, 2018). This suggests that fMRI in humans has untapped potential, and has inspired ongoing efforts into increasing the imaging resolution of MRI in order to take full advantage of the intrinsically high precision of blood flow regulation (Feinberg et al., 2023). Alongside efforts into advancing data acquisition, parallel efforts into improving fMRI data analysis to take into account what is currently known about the hemodynamic response and how it is shaped by the local vascular anatomy open a promising avenue to help enhance the interpretation of these data to achieve greater neuronal specificity (Polimeni et al., 2018). Such “anatomically-informed” analysis strategies may help extract more meaningful information about human brain function.

Of all fMRI contrasts, the blood-oxygenation-level-dependent (BOLD) contrast, which reflects changes in the deoxyhemoglobin content of blood, is the most commonly used due to its simplicity, robustness and high sensitivity, yet it is well known to exhibit poor specificity. The strongest BOLD-fMRI responses are found in large draining veins that are far from the site of neuronal activity. In the cerebral cortex, the BOLD-fMRI response is known to initiate within the tissue parenchyma microvasculature and then spread in time to the downstream draining veins; deoxygenated venous blood collects upwards to the cortical surface into the large pial vein in approximately 0.5 s, which is observable in human fMRI and has been reported in high-resolution small animal work (Yu et al., 2012). In this way, hemodynamics are shaped by the anatomy and physiology of vascular supply and drainage across scales (Hillman, 2014). Because any given pial vein will drain deoxygenated blood from large cortical domains, the large BOLD signals seen in these veins are spatially non-specific (Jonathan R. Polimeni et al., 2010). However, the earliest stages of the BOLD response will likely be closer to the site of neuronal activity (Blazejewska et al., 2018; Sirotin et al., 2009) and thus are hypothesized to exhibit improved specificity in both space and time.

These vascular influences on the spatiotemporal evolution of the hemodynamic response are especially relevant for “laminar fMRI” studies that seek to localize activity to individual cerebral cortical layers (Norris and Polimeni, 2019). In this context, a more complete understanding of the hemodynamic response timing would allow investigators to identify the first cortical layer within a given cortical area to activate—this would provide the most straightforward means to identify the input layer along the feedforward or feedback pathway that is engaged by the particular stimulus or task (Petridou and Siero, 2019a), which can be used decipher functional circuitry in the human brain (Huber et al., 2021; Jonathan R Polimeni et al., 2010). The patterns of BOLD-fMRI responses across the cortical thickness will then evolve in time, such that as the response spreads it will increasingly reflect not just the neuronal activity of interest but also the influences of the vascular architecture and the microvascular hierarchy of the cortex, complicating the interpretation of the hemodynamic response in terms of the underlying neuronal activity.

Indeed, previous human fMRI studies have demonstrated differences in the BOLD-fMRI responses across cortical depth, reflecting spatial organization of the vascular architecture, with smaller-amplitude responses seen within the parenchyma near the microvasculature and larger-amplitude responses seen at the pial surface near to large draining veins (Koopmans et al., 2010; Jonathan R. Polimeni et al., 2010). It is now common to assume that, because large draining veins are found on the cortical surface, the BOLD-fMRI response should steadily increase in amplitude as one progressively samples the BOLD response closer to the pial surface. However, this simplifying assumption may not hold at every cortical location—vascular architecture varies somewhat from location to location in the cortex, and large pial veins are somewhat sparse (Bollmann et al., 2022; Duvernoy et al., 1981), therefore a strict monotonically-increasing BOLD response amplitude from parenchyma to the pial surface may not be universal. In fact, studies have suggested substantial and perhaps meaningful spatial heterogeneity of cortical-depth profiles of gradient-echo BOLD (Fracasso et al., 2018a; Gomez et al., 2022; Jonathan R. Polimeni et al., 2010), with the highest spatial variability of the profiles observed near the pial surface (Jonathan R. Polimeni et al., 2010).

Laminar fMRI, which employs sub-millimeter imaging resolution, typically performs spatial averaging of cortical-depth profiles inside a specific region-of-interest (ROI) in the targeted cortical area, assuming similar cortical-depth profiles at each location within the ROI. This pooling reduces biases associated with sampling some depths more than others, which happens due to the limited resolution of fMRI and the folding of the cortex (Koopmans et al., 2011; Polimeni et al., 2018) and increases SNR, thus enabling small-voxel laminar-fMRI experiments—even at lower (e.g., 1.5T) magnetic field strengths (Markuerkiaga et al., 2021). The gain in SNR also allows the use of fMRI contrasts with higher spatial specificity but lower functional sensitivity than BOLD (Huber et al., 2019). Therefore, testing the assumption of similar cortical depth profiles at each location within the ROI is crucial for laminar fMRI from the methodological perspective.

Motivated by this, in this work we investigate temporal and spatial characteristics of cortical-depth response profiles in the human visual cortex and test the hypothesis that the temporal features of the BOLD-fMRI response varied over space, and whether spatial features varied over time. We used a common visual stimulus and measured responses within human primary visual cortex (V1) where the vascular architecture is well known. In testing variability over space, we considered both *across-depth* variations of the BOLD response (cortical-depth profiles) and *across-region* variations within V1. In testing variability over time, we analyzed evolution of cortical-depth profiles as a function of post-stimulus time. We applied k-means clustering at various time-points to identify cortical locations exhibiting different depth profiles. Finally, we performed a series of analyses aiming to test whether the differences between the profiles are meaningful, not due to sampling biases caused by the anatomical geometry or dominated by noise in the data. Not only does our approach test whether the local vascular architecture and the known temporal evolution of the hemodynamic response yield the expected spatiotemporal spread of the BOLD-fMRI signal, but our findings may also inform methodological approaches aiming to improve spatio-temporal localization of the neuronal component of BOLD fMRI signal—thereby enhancing neuronal specificity of the measurements by accounting for the influences of the vascular anatomy and physiology.

## METHODS

### Subjects and data acquisition

In this study we perform single-subject analyses and investigate hemodynamics in individual participants. Therefore we recruited six healthy volunteers (3M/3F, 28±8y.o.) who provided prior written informed consent, in accordance with our institution’s Human Subjects Research Committee policies. Imaging was performed on a whole-body 7T scanner (MAGNETOM Terra, Siemens Healthineers, Erlangen, Germany) equipped with an inhouse-built 64-channel receive-coil brain array (Mareyam et al., 2019). In each experimental session, we acquired a 0.75-mm isotropic resolution FOCI-MEMPRAGE data (Hurley et al., 2010; van der Kouwe et al., 2008; Zaretskaya et al., 2018) and a standard two-echo gradient-echo B_0_ field map. For functional scans, we acquired 10 BOLD-weighted fMRI runs per session, which were positioned on the calcarine sulcus and oriented approximately coronally with the slice normal roughly parallel to the long axis of the calcarine, using standard 2D-gradient-echo EPI protocols at 0.8-mm isotropic resolution. (See **Table 1** for detailed acquisition parameters.)

**Table 1.** Summary of scanning parameter values. Repetition time (TR), echo time (TE), inversion time (TI), field of view (FOV), number of slices (SL), resolution (RES), flip angle (FA), bandwidth (BW), nominal echo spacing (ESP), in-plane acceleration factor (*R*), number of time points (TP), and acquisition time (TA).

| sequence | TR [ms] | TE(s) [ms] | TI [ms] | FOV [mm] | SL | RES [mm] | FA [°] | BW [Hz/px] | ESP [ms] | R | TP | TA |
| --- | --- | --- | --- | --- | --- | --- | --- | --- | --- | --- | --- | --- |
| FOCI-MEMPRAGE | 2530 | 1.8 3.2, 4.7, 6.2 | 1100 | 240 x 240 | 224 | 0.75 | 6.6 | 1300 | 8.10 | 2 | --- | 7:23 |
| 2D GE EPI | 2000 | 27 | --- | 192 x 192 | 31 | 0.80 | 74 | 1190 | 1.01 | 3 | 140 | 5:03 |
| 2D GE $B_0$ map | 1000 | 3.94, 4.96 | --- | 224 x 224 | 32 | 1.6 | 35 | 1553 | | --- | --- | 4:44 |

During each functional scan, subjects were presented with a standard visual stimulus: a full-field-of-view black-and-white ‘scaled-noise’ pattern, counter-phase flickering at 8 Hz, in four randomly-jittered blocks of 16-s duration and 40–49-s inter-stimulus-intervals. This resulted in four stimulus trials per run, and 10 runs per session, for a total of 40 trials per session.

### Data processing

We performed combined bias-field correction and tissue segmentation of the T_1_-weighted anatomical data using the unified segmentation approach implemented in SPM (https://www.fil.ion.ucl.ac.uk/spm/), (Ashburner and Friston, 2005). Surface-mesh representations of the cerebral cortical gray matter including the gray-white and gray-CSF interfaces were reconstructed automatically using FreeSurfer version 7.1.1 (https://surfer.nmr.mgh.harvard.edu/), (Fischl, 2012) based on brain and white matter (WM) segmentations. These segmentations were then manually corrected based on tissue probability maps previously obtained from SPM, after which the surface-reconstructions were repositioned (see **Figure 1a**). Additional surface-meshes were then generated corresponding to cortical depths spaced uniformly every 10% between WM and pial-surface to provide an “equi-distant” sampling across depths (Polimeni et al., 2018).

**Figure 1.**
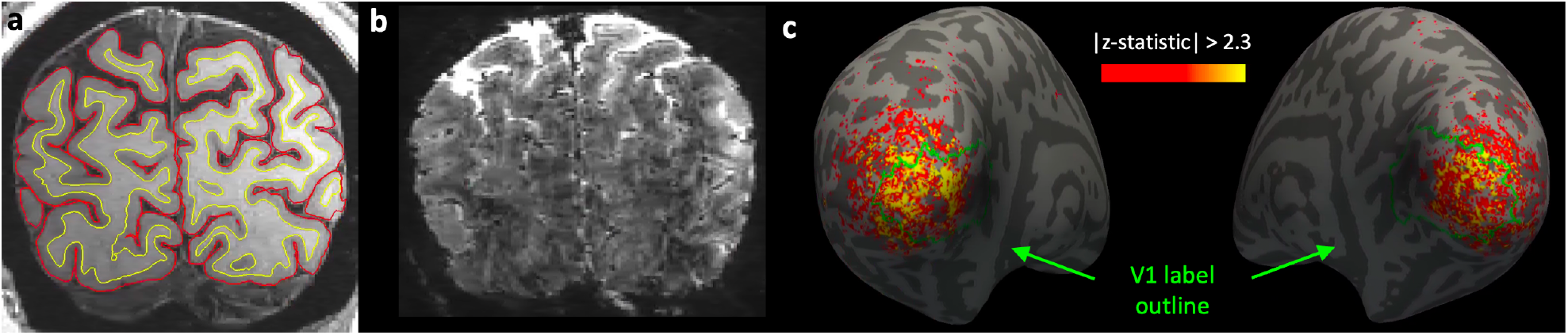
Example data for a representative subject. **(a)** A coronal cross-section of the MEMPRAGE, root-mean-square (RMS) combination of the images acquired with four echo times (see Table 1), with overlaid white matter (yellow) and pial (red) surface contours representing the intersection of the cortical surface reconstruction. **(b)** A coronal slice from an average volume of the first 5 frames (representing baseline) of a single fMRI run, previously corrected for motion and linear trends. **(c)** Inflated surfaces of left and right cerebral hemispheres, with overlaid z-statistic values combined across all runs (using a fixed-effects analysis) and thresholded to |z|>2.3; an automatically-generated anatomical prediction of V1 based on the cortical folding pattern (outlined in green) was used to create an intersection with the activation region-of-interest (ROI).

EPI data were motion corrected using 3dvolreg from the AFNI toolbox (https://afni.nimh.nih.gov/) and linear trends were removed from the time-series data. **Figure 1b** shows an example coronal slice from an average volume of the first 5 frames (baseline) of a single preprocessed fMRI run. In order to co-register our EPI-based BOLD-weighted fMRI data with our T_1_-weighted anatomical data, boundary-based registration (Greve and Fischl, 2009), initialized with manual alignment, was performed in two steps: first based on aligning cortex across the whole-brain excluding regions prone to large susceptibility-induced EPI distortions identified based on FreeSurfer labels^1^, which was then refined in a second step constrained to only align cortex within the FreeSurfer V1-mask.

To define regions-of-interest (ROIs), a standard general linear model (GLM) analysis was performed in volume space using the feat tool provided by FSL (https://fsl.fmrib.ox.ac.uk/fsl). The resulting z-statistic maps were combined across all runs a using fixed-effects analysis, thresholded (*z* > 2.3) and projected onto the cortical surface-meshes corresponding to WM/GM interface. ROIs were then defined based on the activation at WM/GM interface to avoid pial vein bias influence. The final non-contiguous ROIs, one per hemisphere, were generated by intersecting these initial, functional ROIs with anatomical V1 labels (automatically generated by FreeSurfer) to limit the analysis to locations within the primary visual cortex (see **Figure 1c**).

Dynamic activation maps calculated at every post-stimulus time-point were created for each fMRI run then combined across runs, similar to our previous study (Blazejewska et al., 2019a). These so-called dynamic statistical parameter maps (dSPM) (Dale et al., 2000) consisting of t-statistic values were calculated as signal change (Δ*S*) divided by the temporal standard deviation of the residuals (σ). The transformations obtained from the registration of the fMRI data to the anatomical reference data were applied to project dSPM maps onto the surface-meshes from which one cortical depth profile of dSPMs can be obtained for each vertex (across 11 depths) of the surface mesh.

### Evolution of BOLD-fMRI response in time and across cortical depth

The dSPM values were averaged within the functionally-defined ROIs, combined across both hemispheres, averaged across all subjects. To summarize how the activation varied with both time and cortical-depth, 2D “time-depth map” was generated, where t-statistic values were plotted across time (from 1 to 30 s post-stimulus onset) along the horizontal axis and across cortical-depth (from white matter to pial surface) along the vertical axis.

### Clustering of cortical-depth response profiles

To identify cortical locations that exhibited similar cortical depth activation profiles, a K-Means Clustering approach was used. For each subject, the analysis was applied to the cortical-depth profiles of the t-statistics generated from the dSPM analysis, restricted to vertices within the activated ROI (combined across the two cortical hemispheres), at each time-point ranging from 6 to 22 s post-stimulus (every 1 s). The cost function used for clustering, which defined similar cortical-depth profiles, was a simple root-sum-of-squares difference calculated between the 11 sampled cortical depths (or the “Euclidean distance” in this 11-dimensional space) to account for both, differences in response profile shape and amplitude across cortical locations. Several different numbers of clusters, *k*, were tested (ranging from 2 to 6) and *k*=4 was selected because it provided the smallest number of activation profiles that clearly differed by eye, hence yielding one distinct cortical-depth response profile per cluster. This generated *k*=4 cluster ‘centroids’, which represent the typical cortical-depth profile for each cluster (calculated as mean of all profiles in a given cluster), and a set of cluster-membership labels that assigned to each cortical location a single cluster label (1 to 4). To then test whether each cluster exhibited a distinct evolution across both time and cortical-depth, we generated a “time-depth map” for each cluster and compared them qualitatively. To then more closely assess the consistency of activation within each cluster as a function of post-stimulus time, we compared the plots of cortical-depth profiles for each cluster at several post-stimulus time-points. To investigate which components of t-statistic (i.e., activation effect size or the noise variance) might be driving the differences between the cortical-depth profile shapes, and to inspect the consistency of these contributors across all ten runs, signal baseline (*S*_0_) calculated as median of the signal before the first task block for a given run, percent signal change (Δ*S*/*S*_0_) and temporal standard deviation of the residuals (σ) were compared across clusters.

We considered that the observed differences in the shapes of the trial-averaged BOLD response profiles between clusters may result simply from biases in the way that the fMRI data are irregularly sampled by the cortical-surface meshes and interpolated onto the mesh vertices: either due to the varying geometry of the cortical surface or due to local geometric distortion of the EPI data. To test for these possible artifactual, systematic relationships between features of the cortical geometry and cluster membership that might suggest sampling/interpolation biases, histograms of cortical thickness and cortical curvature of all locations within each cluster were plotted. Additionally, to test for sampling biases due to EPI distortions, B_0_ field offsets were calculated from separately acquired B_0_ field maps and histograms were calculated again for all locations within each cluster.

Consistency of the clusters within each subject was investigated with simple train-test cross-validation approach: five out of ten runs were selected randomly, and clustering was performed on average t-statistic calculated for this subset. Then profiles of average t-statistic across the remaining five runs were plotted within these clusters for testing.

### Comparing clustering results to ‘null’ profiles generated from random noise

It is mathematically possible that k-means clustering of random noise could result in a set of smooth, mutually-orthogonal cortical-depth profiles (analogous to a “partition of unity”) that would not meaningfully capture the typical profile within any cluster. If clustered profiles, on average, exhibited a trend increasing towards the pial surface, with a superimposed high-level of statistically independent random noise, we may expect to see a similar set of cortical-depth profiles to those observed from our fMRI data clustering approach. Therefore, to test whether clusters with cortical-depth profiles similar to those found in our data could be generated from simply clustering random noise, we performed two experiments analyzing synthetic data generated from random noise to establish what characteristics would be seen in ‘null ‘cortical-depth profiles.

First, we generated a synthetic gaussian-distributed noise dataset consisting of random cortical-depth profiles, with mean and standard deviation matched to the spatial mean and spatial standard deviation of our t-statistic cortical-depth profiles (within the activated ROI) calculated from one example subject. These synthetic-noise cortical-depth profiles were then clustered following identical procedures as for the actual fMRI data. The cortical-depth response profile centroids and corresponding cluster-membership labels were generated, as well as histograms of the distance from the cluster centroid for each cluster.

Second, to further ensure that the clusters are not simply a result of spatially-structured smoothing of our high-resolution fMRI data imposed by interpolation during motion correction (Polimeni et al., 2018; Wang et al., 2022), we generated 4D synthetic-noise datasets (three spatial dimensions plus one temporal dimension) matching the dimensions of our acquired BOLD-weighted EPI data. Each 4D synthetic-noise dataset consisted of i.i.d., spatially and temporally white gaussian random noise, and one such 4D synthetic-noise dataset was created for each of the 10 runs for an example subject. Motion correction based on the motion parameter estimates calculated from the measured fMRI data was applied to each corresponding run of 4D synthetic-noise data, which imposed identical smoothing due to interpolation on the 4D noise data that was applied to our fMRI data during preprocessing. Each of the motion-corrected fMRI data runs for this subject as well as the corresponding, preprocessed 4D synthetic noise were clustered. Furthermore, cluster membership obtained from the fMRI data was then applied to the preprocessed 4D synthetic-noise and vice versa on a per-run basis, and the resulting cortical-depth profiles averaged within each cluster were plotted. In addition, run-to-run cross-validation was performed for both the fMRI data and preprocessed 4D synthetic-noise data, where clusters from an arbitrarily selected run were applied to another run, and again the corresponding cortical-depth profiles were plotted.

## RESULTS

### Evolution of BOLD-fMRI response across time and across cortical depth

For each subject, analysis was performed inside the activated ROI which included 2502±888 cortical locations and corresponding to them cortical-depth response profiles (see **Figure 1c**). **Figure 2** summarizes the evolution of the trial-averaged BOLD-fMRI responses in time post-stimulus onset and across cortical depth, averaged over all six subjects. The cortical-depth profiles of the BOLD response plotted for selected time-points between 6 and 22 s post-stimulus onset demonstrate that, as predicted, the BOLD response at early time-points is maximal within the parenchyma, and only becomes maximal at the pial surface at

**Figure 2.**
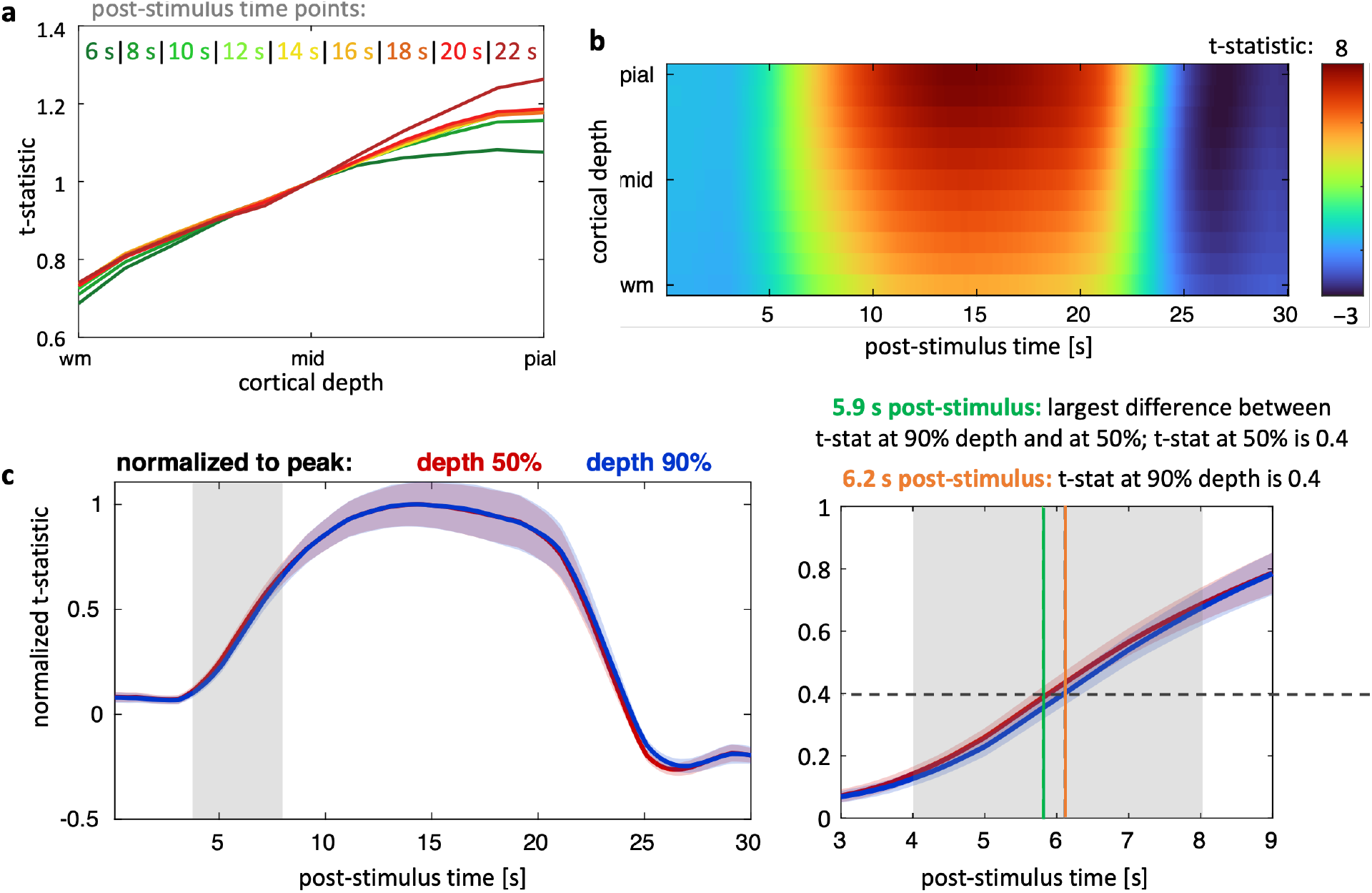
Evolution of BOLD fMRI response in time & across cortical depth, average across all subjects. **(a)** Cortical-depth response profiles plotted for selected time points 6–22 s post-stimulus onset, normalized to the response measured at mid-depth for each time point. **(b)** 2D t-statistic map, cortical depth × post-stimulus time. **(c)** Response evolution in time for the first 30 s post-stimulus onset plotted for two selected cortical depths: mid-depth (50%) and near-pial (90%) (shaded areas represent error across subjects), with a magnified section (right) where the responses for these two depths differ (between 3 and 8 s post-stimulus, marked with gray shading).

later time-points (**Figure 2a**). To visualize how the trial-averaged BOLD response changes over both time and space, and the interaction between these two dimensions, we generated 2D time-depth plots of the t-statistic computed for each post-stimulus time and cortical depth, presented in **Figure 2b**. These plots show the lack of separability between the two dimensions (time and space) and that, indeed, the BOLD response evolves over time differently at each cortical depth. Comparison of the evolution of the BOLD responses at mid-depth (i.e., 50% depth) and near-pial-depth (i.e., 90% depth) showed that, at early time points (until about 8 s post-stimulus) the response at mid-depth is on average about 0.3 s faster than that of the near-pial-depth (**Figure 2c**).

### Spatial heterogeneity of cortical-depth response profiles

The results of k-means clustering of the t-statistic cortical-depth profiles measured 8 s post-stimulus onset at each activated cortical location demonstrated spatial heterogeneity of cortical-depth profiles of BOLD activation across V1, as presented in **Figure 3** for a representative subject. Four cortical-depth profile shapes were identified, as shown in **Figure 3a**: Cluster #1 having a relatively flat profile (green, ∼37% of vertices), Cluster #2 having a profile that is maximal at mid-depth (purple, ∼25% of vertices), Cluster #3 having a profile that increases approximately monotonically from the white to the pial surface (red, ∼28% of vertices) and Cluster #4 a profile with a high amplitude that is maximal in near-pial depth (orange, ∼10% of vertices). The corresponding 2D time-depth plots for each cluster are shown in **Figure 3b** (similar to **Figure 2b**), where earlier response at mid-depths is especially apparent in a cluster without pial bias (Cluster #2, purple). All cluster profiles differed markedly from the average profile corresponding to the spatial average across the entire activated ROI (**Figure 3c**).

**Figure 3.**
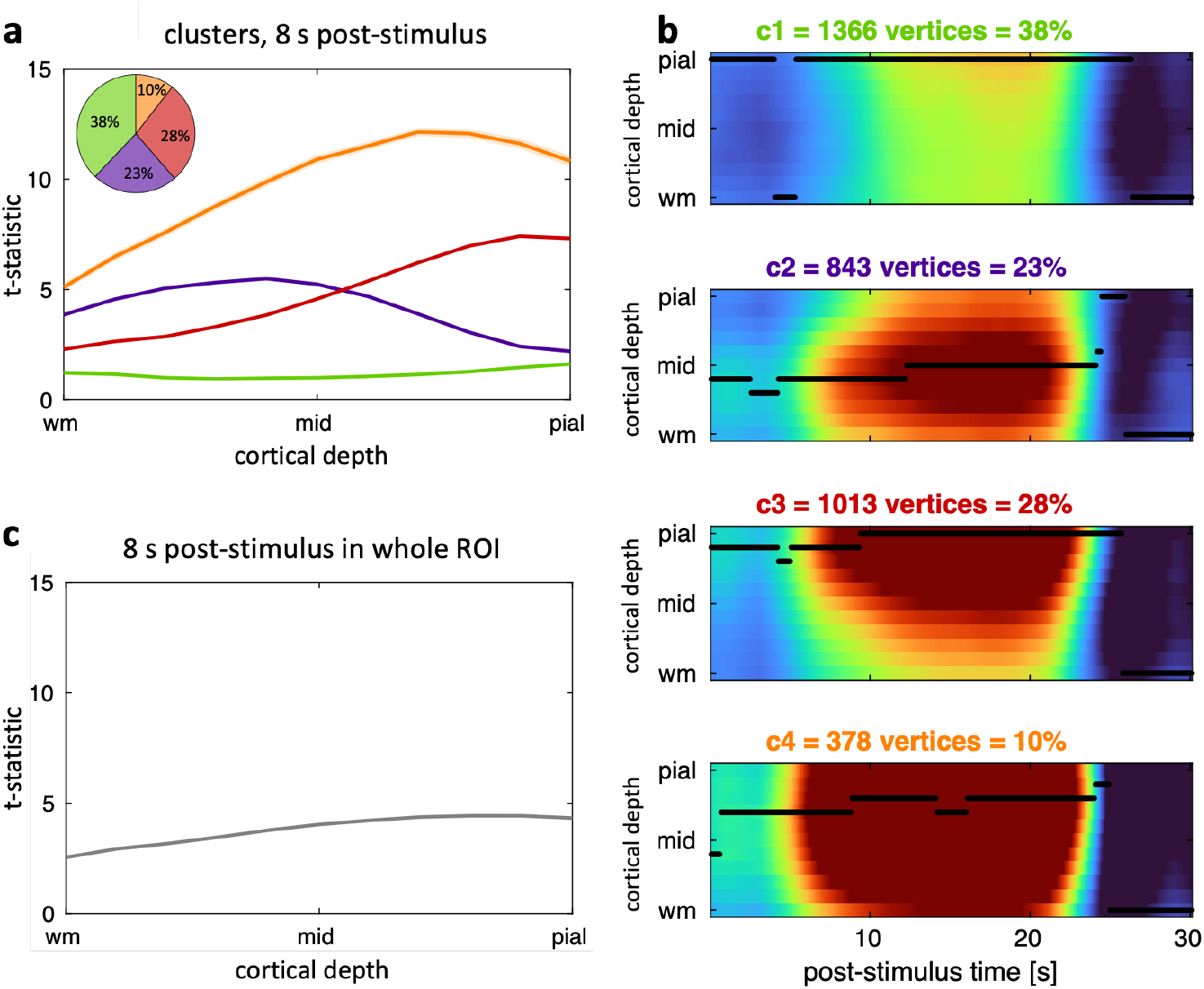
Results of K-Means Clustering of cortical-depth profiles of BOLD-fMRI responses 8 s post-stimulus onset for a representative subject. **(a)** Average cortical-depth response profiles within the four clusters 8 s post-stimulus onset, and pie chart of the percentages of activated vertices in each cluster. **(b)** The 2D time-depth maps of t-statistic values for each cluster; the 95^th^ percentile is indicated by the black line and is included to indicate the cortical depth corresponding to the maximal response amplitude at each time point. **(c)** Average cortical-depth response profile 8 s post-stimulus onset across the entire activated ROI for reference.

To confirm the stability of the clusters over time, and test whether they may reflect fixed anatomical features, we re-computed the clusters at different post-stimulus time-points between 6 s (early response) and 20 s (late response) post-stimulus onset, and compared the shapes of the resulting cortical-depth profiles as well as the numbers of vertices within each cluster and the spatial arrangement and organization of the clusters. The results are presented in **Figure 4**, which demonstrates that the basic shapes of the cortical-depth profiles for each cluster were largely consistent over time, with small changes in amplitude seen at different post-stimulus times, as can be seen also in the plots of **Figure 2**. While the exact percentages of vertices belonging to each cluster varied slightly when the clustering was performed at different post-stimulus time points, the spatial arrangement of the clusters was stable, indicating that cluster membership appears fixed over time and thus it is likely that the clusters reflect some aspect of the functional/vascular anatomy that does not change with time.

**Figure 4.**
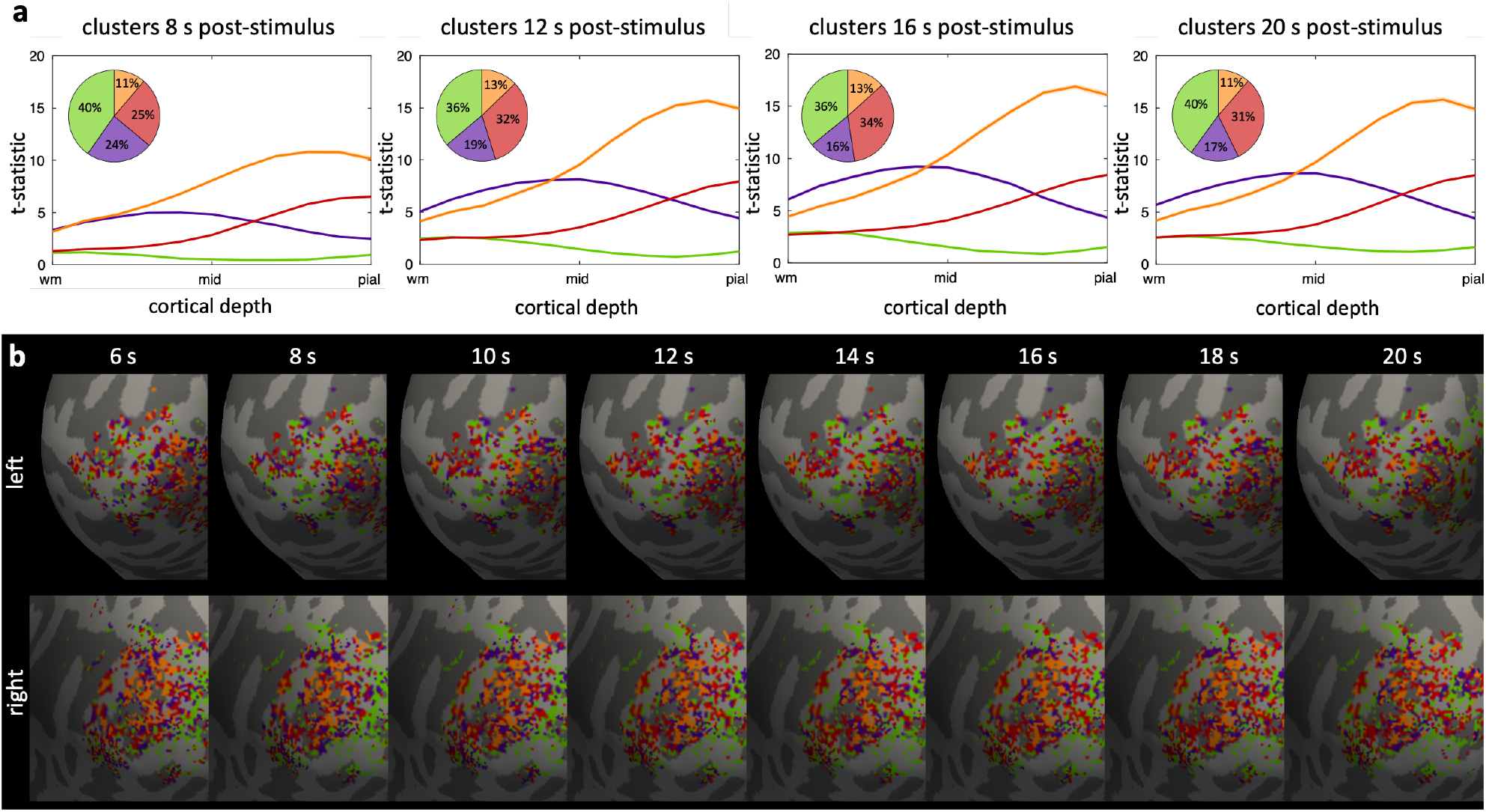
K-means clustering of cortical-depth BOLD fMRI response profiles at different time-points post-stimulus onset for a representative subject. **(a)** Average cortical-depth response profiles inside 4 clusters 8, 12, 16 (peak) and 20 s post-stimulus onset. **(b)** Color-coded labels representing clusters computed from data sampled at different time-points 6–20 s post-stimulus onset overlaid onto inflated cortical surface of left and right hemisphere.

We also tested the consistency of the clusters over time by performing clustering on data sampled at one post-stimulus time point and then applying these cluster definitions to data sampled at another post-stimulus time point. In other words, we assigned each cortical location or each surface vertex to one of the four clusters based on computing the k-means clustering of the trial-averaged BOLD response sampled at one post-stimulus time point, then compared the cortical-depth profiles of the BOLD response within these clusters both using the BOLD response at this time point and the BOLD response at another time point. **Figure 5a & b** shows the cortical-depth profiles for each cluster defined based on the BOLD response at 8 s post-stimulus for the BOLD response at 8 s post-stimulus as well as the BOLD response 16 s post-stimulus (which roughly corresponds to the time at which the BOLD response amplitude peaks). We see again remarkable consistency between clusters derived using these two time points, again suggesting that the cluster membership is stable and reflects some fixed features of the functional/vascular anatomy.

**Figure 5.**
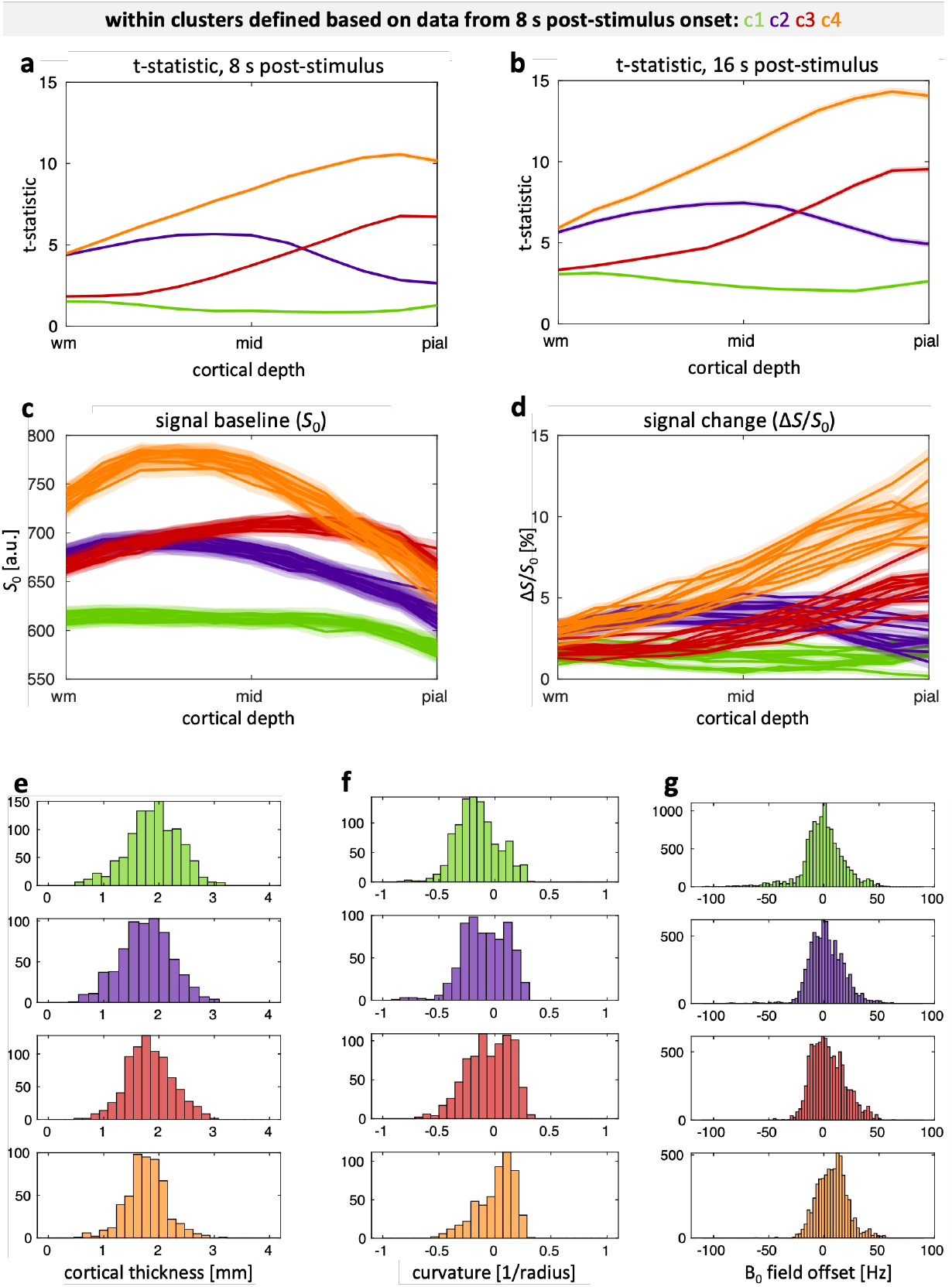
Validation: clusters of cortical-depth profiles 8 s post-stimulus onset, for a representative subject. **(a)** Average cortical-depth profiles of BOLD response within the clusters defined from the data sampled at 8 s post-stimulus onset. **(b)** Average cortical-depth response profiles inside the clusters 16 s post-stimulus onset (response peak). **(c)** Cortical-depth profiles of signal baseline inside the clusters for each of ten runs. **(d)** Cortical-depth profiles of percent signal change inside the clusters for each of ten runs; **(e)** Histograms of cortical thickness. **(f)** Histograms of cortical curvature. **(g)** Histograms of B_0_ field offset, inside the four clusters.

To gain insight into what anatomical features may be determining the different cortical-depth profiles of the BOLD response across clusters, we performed similar comparisons by plotting different features of the fMRI data including the signal baseline (*S*_0_) and percent signal change of the trial-averaged BOLD response at specific post-stimulus time points (Δ*S*/*S*_0_). The cortical-depth profiles of signal baseline (**Figure 5c**) and percent signal change (**Figure 5d**) were both distinct across the four clusters—again, clusters derived from the BOLD response t-statistic—indicating that cluster membership derived from one aspect of the fMRI data generalizes to other aspects of the fMRI data. The cortical-depth profiles of signal baseline were highest in Cluster #4 (**Figure 5c**, orange) and lowest in Cluster #1 (**Figure 5c**, green), which somewhat matches the corresponding profiles of the BOLD response t-statistic, however the profile shapes were otherwise different between signal baseline and BOLD response t-statistic, indicating that the underlying anatomical features that drive cluster membership have different effects on these two aspects of the fMRI data. Notably, the cortical-depth profiles of the signal baseline all exhibited a minimum at the pial surface, and a maximum within the parenchyma, unlike the cortical-depth profiles of the BOLD response. The cortical-depth profiles of percent signal change, however, more closely resembled the t-statistic profiles, as expected because of the similarity of these two measures of activation amplitude. While both t-statistic and percent signal change measure the BOLD effect, the t-statistic includes a noise or variance term, and so these plots suggest that it is the amplitude of the BOLD response and not this variance term that likely drives cluster membership.

To test whether cluster membership may be driven by sampling biases imparted by varying cortical thickness or cortical curvature or local EPI geometric distortions, all of which influence partial volume effects in the BOLD-fMRI data—or how much cortical gray matter is contained within the EPI voxel—histograms of cortical thickness, cortical mean curvature and local B_0_ field offsets were calculated for each cluster. The results show that these histograms were nearly indistinguishable across all clusters, suggesting that these perhaps less meaningful features of the cortical anatomy are not a major determinant of cluster membership, and that the cluster definitions are unlikely to be driven by any of these sources of systematic sampling bias (**Figure 5e–g**).

Within-subject consistency of the clusters was confirmed using train-and-test approach on random subsets of runs (**Figure 6**). The average cortical-depth response profiles within the clusters were similar in training and testing subsets of the runs (**Figure 6a & e**). This was confirmed by similar 2D histograms of the profiles which appeared minimally more scattered for the testing dataset (**Figure 6c & g**) and similar carpet-plots of the response-profile distances from the cluster centroid (**Figure 6d & h**). These 2D histograms and carpet-plots both indicate that each cluster is comprised of a set of cortical-depth profiles that exhibit shapes that are similar to the average cortical-depth profile for this cluster.

**Figure 6.**
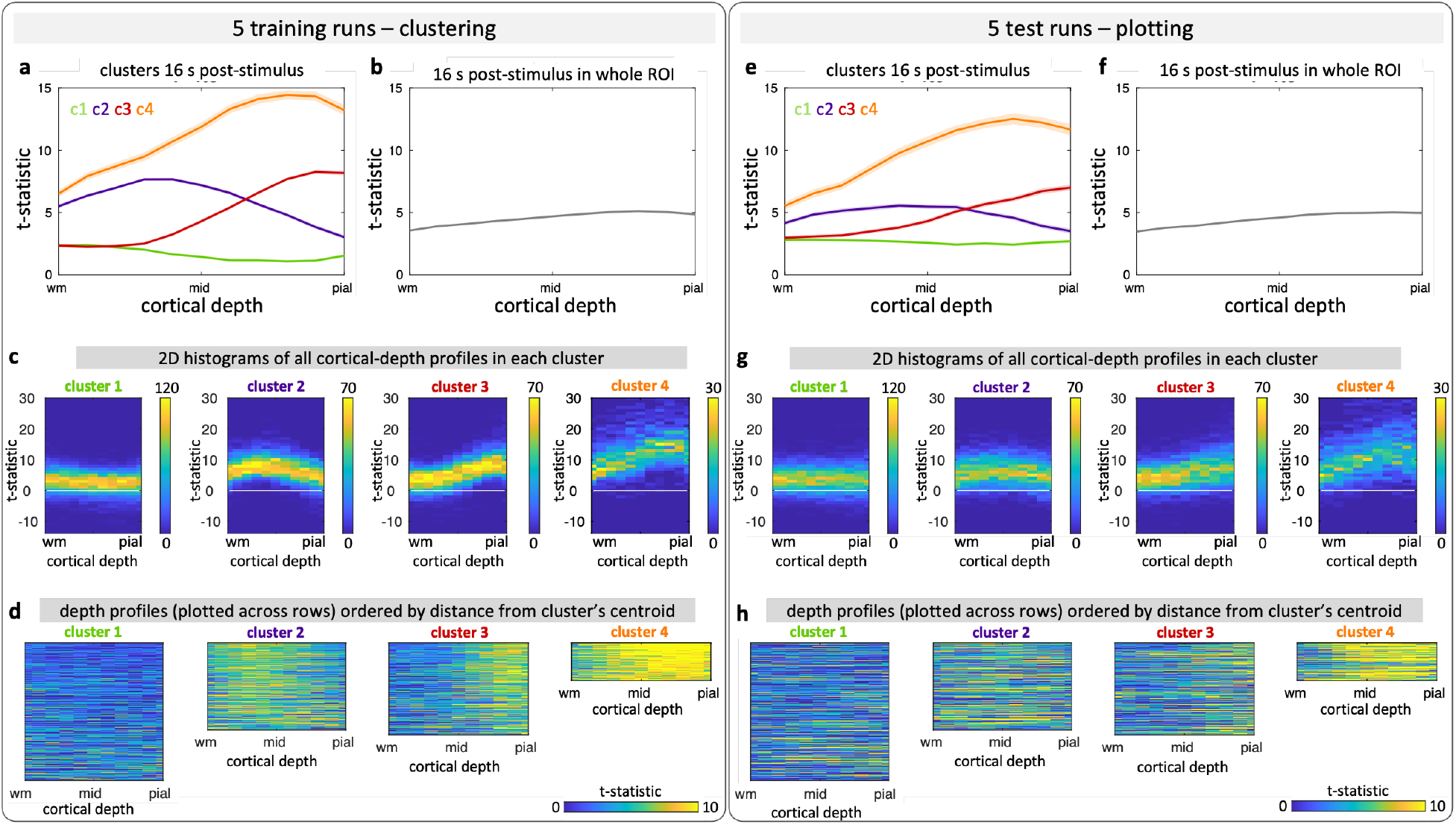
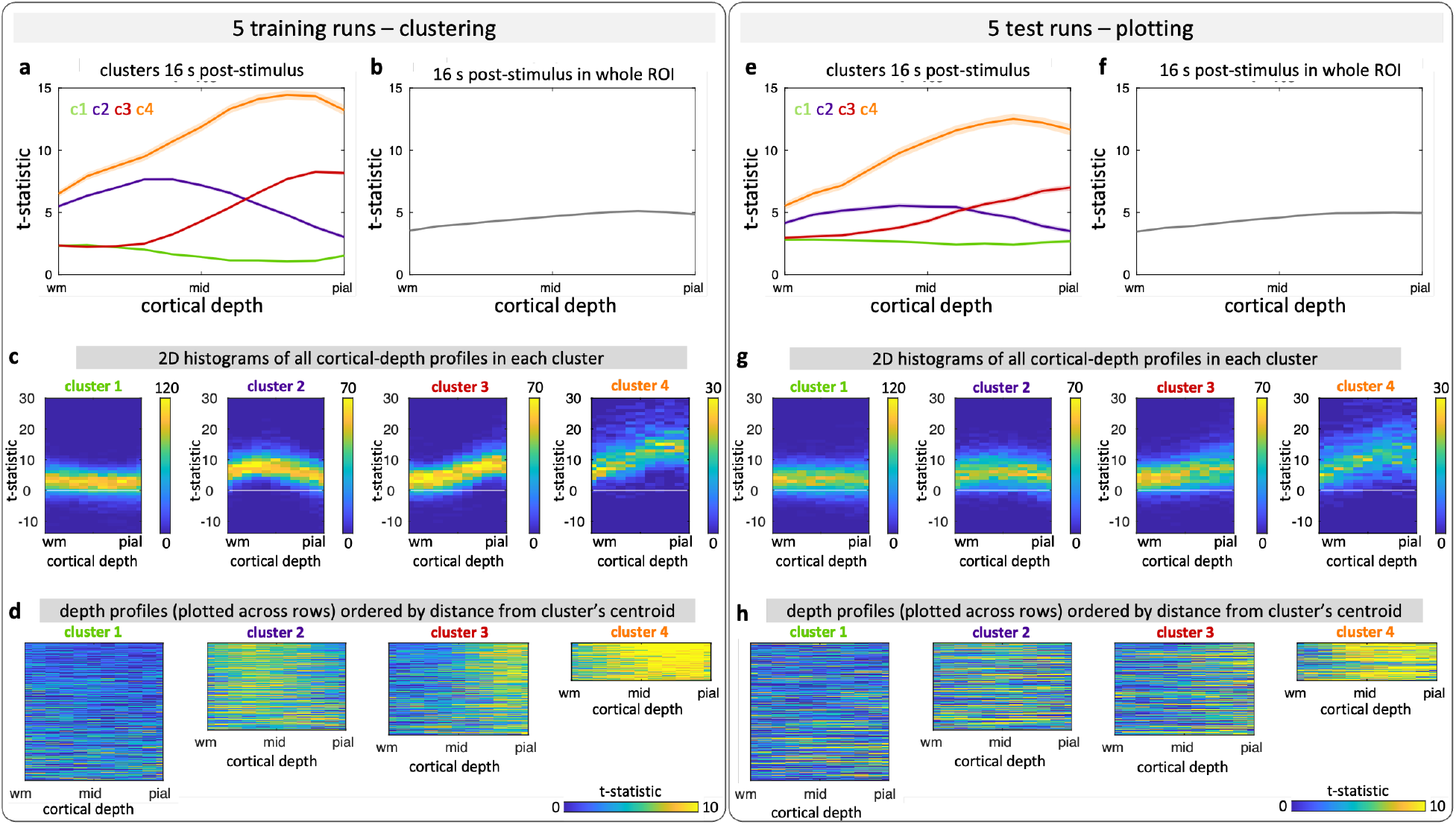
Train/test validation using 5 out of 10 randomly selected runs for training (left) and the remaining 5 runs for testing (right), for a representative subject. Clustering was performed for responses 16 s post-stimulus onset. **(a & e)** Average cortical-depth response profiles inside four clusters. **(b & f)** Average cortical-depth response profile 16 s post-stimulus onset in whole activated ROI. **(c & g)** 2D histograms of response profiles across all runs inside each cluster. **(d & h)** Response profiles (rows) in each cluster ordered (top to bottom) by distance from the cluster’s centroid.

### Clustering of random noise profiles

We found that the clustering algorithm applied to synthetic-noise cortical-depth profiles yield plausible cortical-depth profiles that are similar to those derived from the measured fMRI data (**Figure 7a & d**) but spatial distribution of the vertices within each cluster appeared more structured for the measured fMRI data than for the synthetic-noise profiles (**Figure 7b & e**). In addition, histograms of the response-profile distances from the cluster centroid clearly demonstrated that clusters derived from the measured fMRI data were more compact (shorter distances, narrower distributions) than those derived from the synthetic-noise cortical-depth profiles (longer distances, broader distributions with heavy tails) (**Figure 7c & f**).

**Figure 7.**
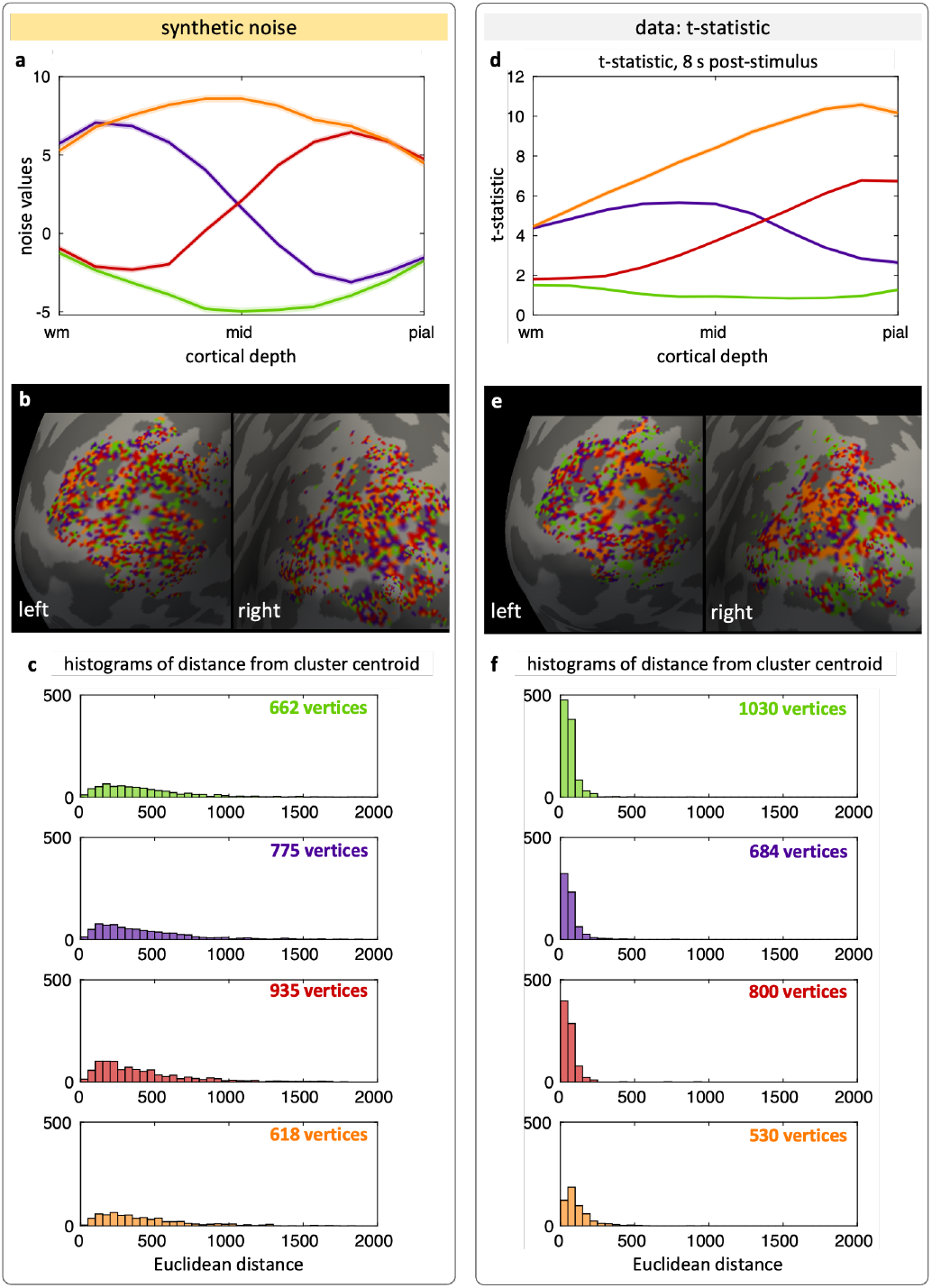
Comparison of k-means clustering results for the BOLD fMRI data and cortical-depth profiles of synthetic, statistically-matched noise. (a & d) Average cortical-depth response profiles of each cluster derived from synthetic cortical-depth profiles (a) and derived from fMRI data (d). **(b & e**) Color-coded labels representing cluster membership overlaid onto inflated cortical surface of left and right hemisphere. **(c & f)** Corresponding histograms of cortical-depth profile distance from each cluster centroid. The histograms of profile distances derived from the fMRI data exhibit a narrow distribution, whereas the histograms of profile distances derived from the synthetic profiles exhibit a heavy-tailed and broad distribution.

Finally, the clusters derived from the preprocessed 4D synthetic-noise dataset yielded inconsistent cortical-depth profiles across separate runs (**Figure 8a & b**), whereas the clusters derived from the preprocessed measured 4D fMRI data were consistent across separate runs (**Figure 8d & e**). As a control analysis, we also applied the cluster-membership labels derived from the preprocessed 4D synthetic-noise dataset to one run of the preprocessed measured 4D fMRI data (**Figure 8c**), which resulted in cortical-depth profiles that were nearly identical across clusters and different from the cortical-depth profiles based on clusters derived from the 4D fMRI data themselves; then, conversely, applied the cluster-membership labels derived from the preprocessed, measured 4D fMRI data to one run of the preprocessed, 4D synthetic-noise dataset (**Figure 8f**), which resulted in cortical-depth profiles that were also nearly identical across clusters and appeared random. Altogether, these analyses demonstrate that the clusters derived from the measured 4D fMRI data are likely not driven by structured smoothing of the data induced by interpolation imposed during data preprocessing.

**Figure 8.**
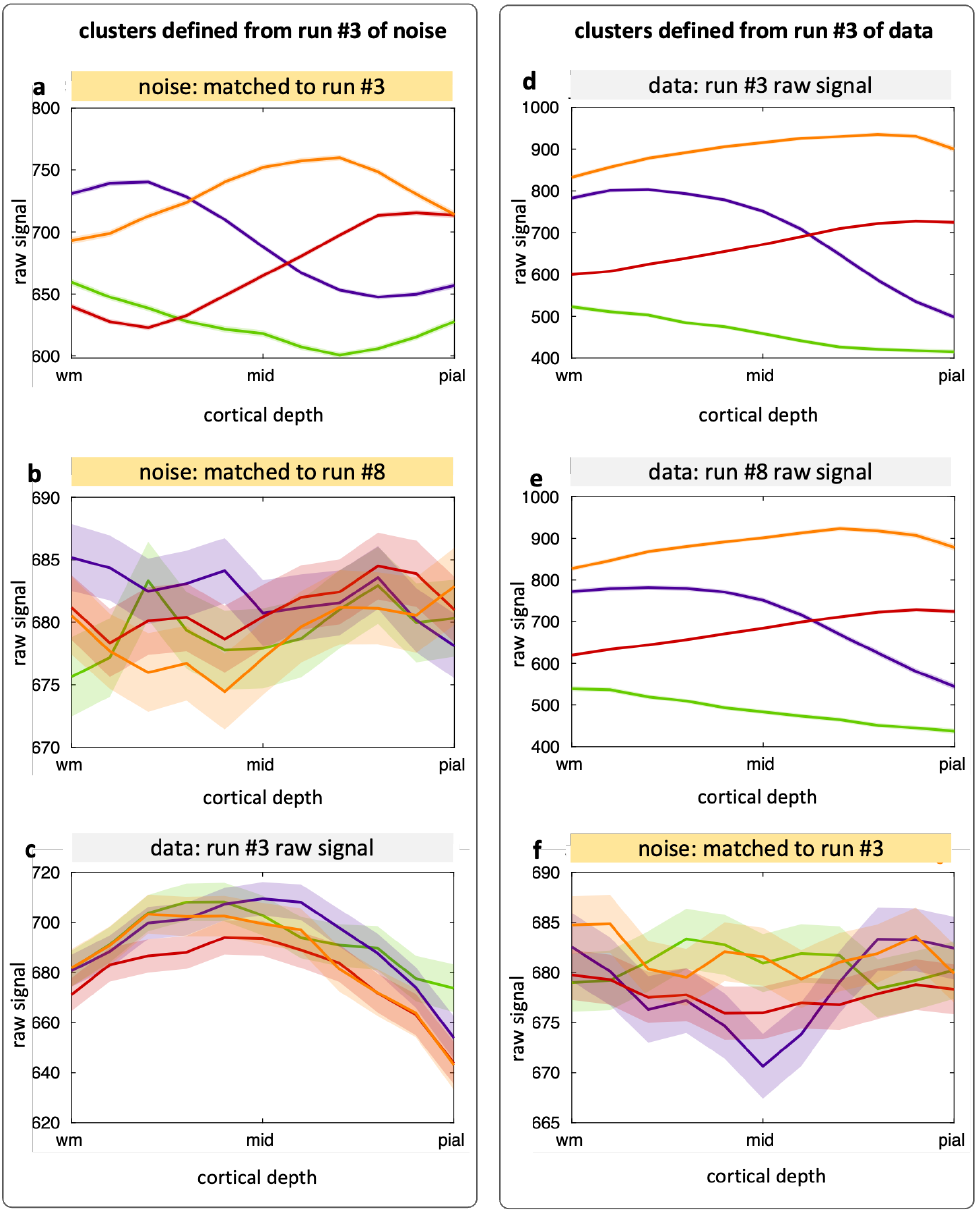
Comparison of k-means clustering results for a representative data and matched gaussian noise after preprocessing. Average cortical-depth response profiles inside the clusters generated for gaussian noise matched to run #3, motion corrected using motion estimates for run #3 (left panel). **(a)** Gaussian noise matched to run #3, motion corrected using motion estimates for run #3. **(b)** Gaussian noise matched to run #8, motion corrected using motion estimates for run #8, **(c)** Motion corrected run #3. Average cortical-depth response profiles inside the clusters generated for a motion corrected run #3 (right panel). **(d)** Motion corrected run #3. **(e)** Motion corrected run #8. **(f)** Gaussian noise matched to run #3, motion corrected using motion estimates for run #3.

## DISCUSSION

We have demonstrated that cortical-depth BOLD-fMRI response-profiles evolve over time and vary substantially over space within the activated region. This suggests that the standard practice of representing the activation across cortical depths by (a) performing a GLM analysis and summarizing the response sampled at any given spatial location over time by a single amplitude or z-statistic or (b) averaging responses as a function of cortical depth and summarizing the response over space by a single cortical-depth profile—or both combined—may result in some loss of meaningful aspects of the underlying fMRI activation patterns.

We found that the early stages of BOLD response ∼6 s post-stimulus onset is initially strongest in mid-cortical depth, and by the time the response reaches its peak amplitude it is strongest at the most superficial depth, at which time the cortical-depth profile exhibits a straight-line trend monotonically increasing towards pial-surface. This result is in agreement with previous high-spatiotemporal resolution studies in small animals (Yu et al., 2012) that also showed that the BOLD response onset appeared earliest in mid-cortical depths. We observed a slight delay between the response measured at mid-depth and the response measured at the pial-surface of about 0.3 s in the early phase of the response (until ∼8 s post-stimulus onset), consistent with animal studies (Tian et al., 2010; Yu et al., 2012) and human studies (Gomez et al., 2024). These results also seem to support the idea that timing of the laminar fMRI responses may provide a tool to detect input layers of the brain circuits as has been previously suggested (Jung et al., 2021; Petridou and Siero, 2019b).

Clustering of the cortical-depth BOLD-fMRI response profiles revealed subsets of the activated vertices with the highest activation localized in mid-cortex appearing at some cortical locations and with the highest activation localized at the pial surface appearing at other cortical locations. The clusters were consistent across different post-stimulus time-points within subject and the average cortical-depth profiles within the four clusters were consistent across subjects. The clustering was independent of cortical features (thickness and curvature) or local geometric distortion that may cause biases in the sampling of the fMRI data, and did not appear driven by structured noise introduced by blurring or interpolation during data preprocessing. Our findings are in agreement with those of previous work where a BOLD response cortical-depth profile “sorting” approach was applied that group profiles into distinct sets, which generated different profile shapes including two monotonic profiles and one with a maximum amplitude at mid-cortical depth (Fracasso et al., 2018b). The spatial distributions of these sets on the cortical surface are similar the spatial distributions our clusters—i.e., exhibiting some spatial structure distinct from random noise—and the differences between sets were attributed to be driven by local vascular anatomy.

It has been shown previously that the gradient-echo BOLD-fMRI signal is strongly influenced by the venous vasculature with the greatest signal changes with activation near large draining veins (Blazejewska et al., 2019a; Olman et al., 2007; Jonathan R. Polimeni et al., 2010). That leads to an average activation profile increasing from WM/GM interface towards the pial surface, resembling a straight-line across cortical depth, as reported previously (De Martino et al., 2013; Huber et al., 2014; Koopmans et al., 2010; Jonathan R. Polimeni et al., 2010). However, due to sparseness of pial vessels on the cortical surface (Bollmann et al., 2022; Duvernoy et al., 1981), it is likely that the cortical-depth profile varies across cortical locations: it is presumably steeper at cortical locations near to large pial veins and shallower in cortical locations far from these large pial veins (Huber et al., 2020). While establishing a direct relationship between cortical-depth response profiles and vascular anatomy was outside of the scope of this work, our results suggest that observed different shapes of the profiles could be due to spatial nonuniformity of the vasculature within the ROI, including both large pial veins and smaller ascending intracortical venules. Our results are also consistent with reports demonstrating marked changes in cortical-depth profiles after excluding locations either with large BOLD activations at the pial surface or near explicitly detected large pial veins, which result in the peak BOLD response shifting from the pial surface to mid-cortical depths (Bause et al., 2020; Chen et al., 2013; Koopmans et al., 2010); if all cortical-depth profiles increased monotonically with depth like the average profile, this removal of fMRI voxels intersecting large pial veins would not result in the observed profile that peaked at mid-cortical depths (see **Supplementary Figure 1**).

**Supplementary Figure 1:**
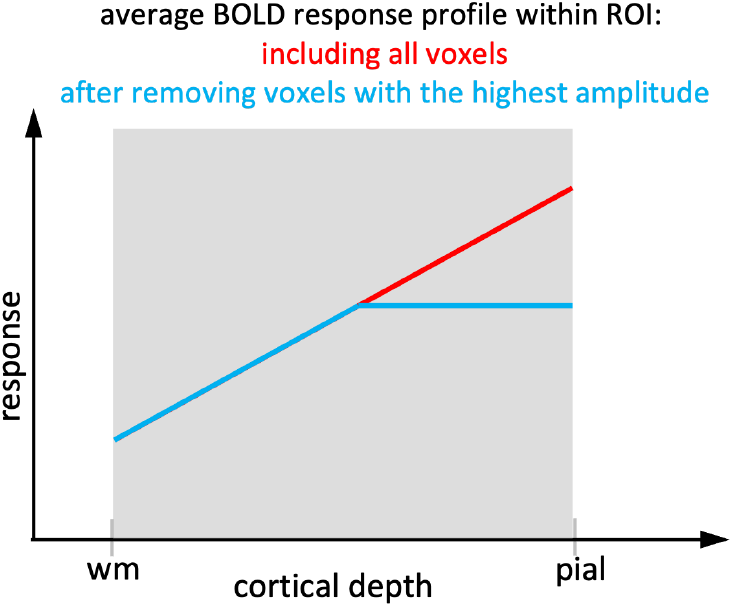
A cartoon illustrating how the average BOLD fMRI response profile shape (plotted with red) would change when voxels with the highest activation are removed (plotted with blue), assuming that all response profiles included in the average have the same shape.

While the observed spatial heterogeneity of the response profiles across the activated locations within V1 could have either vascular or neuronal origins—reflecting either regional differences in vascular architecture or regional differences in the pattern of neuronal activity across cortical layers—the temporal evolution of the measured cortical-depth profiles was on the order of a few seconds, suggesting that this particular aspect is likely explained by a relatively slow hemodynamic effect. In addition, we are not aware of any previous demonstration of such differences in the pattern of neuronal activity across layers within the central visual field representation of V1—with some locations exhibiting a strongest neuronal response to simple visual stimulation in mid-depths and other locations exhibiting a strongest neuronal response in near-pial-depths—again suggesting that the observed spatial variation is also likely attributable to differences in vascular architecture within V1. This includes ‘macrovascular biases’ causing large BOLD responses near pial veins and ‘microvascular biases’ causing local maxima of BOLD responses at cortical depths overlapping with dense capillary beds (Hartung et al., 2023), as well as asymmetries in the timing of the BOLD response caused by deoxygenated blood draining upwards from the parenchyma to the pial surface. However, the lack of direct link between distinct cortical-depth response profiles and the cortical vascular architecture remains a main limitation of our study, and further investigations will be needed to clarify this relationship.

In a broader perspective, our work questions assumptions commonly made in laminar fMRI analysis, when summarizing the activation over time or over space—when applying a standard GLM analysis with a canonical hemodynamic response function for each cortical depth, or when averaging cortical-depth profiles within an ROI—either of which can yield misleading results or cause loss of valuable spatio-temporal information. The implicit assumption that the shape of the cortical-depth profile is constant over all post-stimulus times can potentially result in losing information regarding which layer is activated first. The implicit assumption that that the shape of the cortical-depth profile is constant over all spatial locations within the activated region may similarly lose meaningful layer-specific information since at many locations the BOLD-fMRI response appears to peak in the middle cortical layers and does not exhibit the expected large-vein bias at locations that are presumably between large pial veins and thus not subject to this bias. Based on these observations, alternative approaches to averaging over subsets of activated locations or performing “anatomically-informed smoothing” (Blazejewska et al., 2019b) are promising and could help to eliminate these assumptions and provide enhanced neuronal specificity for gradient-echo BOLD and other fMRI contrasts as well. This preserved information about fine spatial scale and temporal precision of the activation could allow laminar fMRI to detect input/output layers of the within- and across-region brain. In a more general view, alternative “physiologically-informed” fMRI analysis strategies, e.g., employing *microvascular-density weighting* (Chen et al., 2024), could enhance neuronal specificity of fMRI measurements and provide more precise spatio-temporal localization of the underlying neuronal activity in cortical and subcortical brain regions.

In conclusion, we have shown that cortical-depth profiles of BOLD-fMRI responses vary over time and vary over space within the activated region. These variations are likely attributable to biases imposed by vascular anatomy, and these influences of vascular architecture on the spatiotemporal properties of the fMRI response should be carefully considered when analyzing and interpreting laminar fMRI data.

## Acknowledgments

We would like to thank Kyle Droppa for his help with subject recruitment and MRI scanning support, Azma Mareyam for 7T hardware support, and Dr. Saskia Bollmann for valuable discussions. We gratefully acknowledge the use of the Siemens *Works-In-Progress* package WIP#1105.

## Footnotes

1 Regions prone to B_0_-induced EPI distortion that were automatically excluded during boundary-based registration were: middle temporal gyrus, inferior temporal gyrus, temporal pole, fusiform gyrus, entorhinal, medial orbital frontal gyrus, caudal anterior cingulate gyrus, and rostral anterior cingulate gyrus.

## REFERENCES

Ashburner, J., Friston, K.J., 2005. Unified segmentation. Neuroimage 26, 839–851. 10.1016/j.neuroimage.2005.02.018

Bause, J., Polimeni, J.R., Stelzer, J., In, M.H., Ehses, P., Kraemer-Fernandez, P., Aghaeifar, A., Lacosse, E., Pohmann, R., Scheffler, K., 2020. Impact of prospective motion correction, distortion correction methods and large vein bias on the spatial accuracy of cortical laminar fMRI at 9.4 Tesla. Neuroimage 208. 10.1016/j.neuroimage.2019.116434

Blazejewska, A.I., Bernier, M.I., Nasr, S.I., Polimeni, J.R., 2019a. Testing temporal dependence of spatial specificity in BOLD fMRI at 7T: comparing short versus long stimulus duration, in: International Society for Magnetic Resonance in Medicine. p. 0536.

Blazejewska, A.I., Fischl, B., Wald, L.L., Polimeni, J.R., 2019b. Intracortical smoothing of small-voxel fMRI data can provide increased detection power without spatial resolution losses compared to conventional large-voxel fMRI data. Neuroimage 189, 601–614. 10.1016/j.neuroimage.2019.01.054

Blazejewska, A.I., Nasr, S., Polimeni, J.R., 2018. Improved spatial specificity of the early positive BOLD response observed with high-resolution fMRI at 3T, in: International Society for Magnetic Resonance in Medicine. p. 0390. 10.1016/j.neuroimage.2010.09.036.6.

Bollmann, S., Mattern, H., Bernier, M., Robinson, S.D., Park, D., Speck, O., Polimeni, J.R., 2022. Imaging of the pial arterial vasculature of the human brain in vivo using high-resolution 7T time-of-flight angiography. Elife 11, 1–35. 10.7554/eLife.71186

Chen, G., Wang, F., Gore, J.C., Roe, A.W., 2013. Layer-specific BOLD activation in awake monkey V1 revealed by ultra-high spatial resolution functional magnetic resonance imaging. Neuroimage 64, 147–155. 10.1016/j.neuroimage.2012.08.060

Chen, J.E., Blazejewska, A.I., Fan, J., Fultz, N.E., Bruce, R., 2024. Differentiating BOLD and non-BOLD signals in fMRI time series using cross-cortical depth delay patterns * Corresponding author :

Dale, A.M., Liu, A.K., Fischl, B.R., Buckner, R.L., Belliveau, J.W., Lewine, J.D., Halgren, E., 2000. Dynamic Statistical Parametric Mapping: Combining fMRI and MEG High-Resolution Imaging of Cortical Activity. Neuron 26, 55–67.

De Martino, F., Zimmermann, J., Muckli, L., Ugurbil, K., Yacoub, E., Goebel, R., 2013. Cortical Depth Dependent Functional Responses in Humans at 7T: Improved Specificity with 3D GRASE. PLoS One 8, 30–32. 10.1371/journal.pone.0060514

Duvernoy, H.M., Delon, S., Vannson, J.L., 1981. Cortical blood vessels of the human brain. Brain Res. Bull. 7, 519–579. 10.1016/0361-9230(81)90007-1

Feinberg, D.A., Beckett, A.J.S., Vu, A.T., Stockmann, J., Huber, L., Ma, S., Ahn, S., Setsompop, K., Cao, X., Park, S., Liu, C., Wald, L.L., Polimeni, J.R., Mareyam, A., Gruber, B., Stirnberg, R., Liao, C., Yacoub, E., Davids, M., Bell, P., Rummert, E., Koehler, M., Potthast, A., Gonzalez-Insua, I., Stocker, S., Gunamony, S., Dietz, P., 2023. Next-generation MRI scanner designed for ultra-high-resolution human brain imaging at 7 Tesla. Nat. Methods 20, 2048–2057. 10.1038/s41592-023-02068-7

Fischl, B., 2012. FreeSurfer. Neuroimage 62, 774–781. 10.1016/j.neuroimage.2012.01.021

Fracasso, A., Luijten, P.R., Dumoulin, S.O., Petridou, N., 2018a. Laminar imaging of positive and negative BOLD in human visual cortex at 7T. Neuroimage 164, 100–111. 10.1016/j.neuroimage.2017.02.038

Fracasso, A., Luijten, P.R., Dumoulin, S.O., Petridou, N., 2018b. Laminar imaging of positive and negative BOLD in human visual cortex at 7T. Neuroimage 164, 100–111. 10.1016/j.neuroimage.2017.02.038

Gomez, D., Fultz, N., Polimeni, J., Lewis, L., 2022. Temporal properties of fast BOLD fMRI responses in veins and parenchyma, in: International Society for Magnetic Resonance in Medicine. p. 2118.

Gomez, D.E.P., Polimeni, J.R., Lewis, L.D., 2024. The temporal specificity of BOLD fMRI is systematically related to anatomical and vascular features of the human brain. bioRxiv 1–28.

Greve, D.N., Fischl, B., 2009. Accurate and robust brain image alignment using boundary-based registration. Neuroimage 48, 63–72. 10.1016/j.neuroimage.2009.06.060

Hartung, G., Berman, A.J.L., Varadarajan, D., Chen, J.E., Polimeni, J.R., 2023. Computing hemodynamic point-spread functions with biophysical simulations of microvascular networks: effects of trans-laminar capillaries, in: International Society for Magnetic Resonance in Medicine. p. 4027.

Hillman, E.M.C., 2014. Coupling mechanism and significance of the BOLD signal: A status report. Annu. Rev. Neurosci. 37, 161–181. 10.1146/annurev-neuro-071013-014111

Huber, L., Finn, E.S., Chai, Y., Goebel, R., Stirnberg, R., Bandettini, P.A., Poser, B.A., 2021. Layer-dependent functional connectivity methods. Prog. Neurobiol. 207. 10.1016/j.pneurobio.2020.101835

Huber, L., Finn, E.S., Chai, Y., Goebel, R., Stirnberg, R., Stöcker, T., Marrett, S., Uludag, K., Kim, S.G., Han, S.H., Bandettini, P.A., Poser, B.A., 2020. Layer-dependent functional connectivity methods. Prog. Neurobiol. 10.1016/j.pneurobio.2020.101835

Huber, L., Ivanov, D., Krieger, S.N., Streicher, M.N., Mildner, T., Poser, B.A., Harald, E.M., Turner, R., 2014. Slab-Selective, BOLD-Corrected VASO at 7 Tesla Provides Measures of Cerebral Blood Volume Reactivity with High Signal-to-Noise Ratio. Magn. Reson. Med. 72, 137–148. 10.1002/mrm.24916

Huber, L., Uludağ, K., Möller, H.E., 2019. Non-BOLD contrast for laminar fMRI in humans: CBF, CBV, and CMRO2. Neuroimage 197, 742–760. 10.1016/j.neuroimage.2017.07.041

Hurley, A.C., Al-radaideh, A., Bai, L., Aickelin, U., Coxon, R., Glover, P., Gowland, P.A., 2010. Tailored RF Pulse for Magnetization Inversion at Ultrahigh Field. Neuroimage 63, 51–58. 10.1002/mrm.22167

Jung, W.B., Im, G.H., Jiang, H., Kim, S.G., 2021. Early fMRI responses to somatosensory and optogenetic stimulation reflect neural information flow. Proc. Natl. Acad. Sci. U. S. A. 118. 10.1073/pnas.2023265118

Koopmans, P.J., Barth, M., Norris, D.G., 2010. Layer-specific BOLD activation in human V1. Hum. Brain Mapp. 31, 1297–1304. 10.1002/hbm.20936

Koopmans, P.J., Barth, M., Orzada, S., Norris, D.G., 2011. Multi-echo fMRI of the cortical laminae in humans at 7 T. Neuroimage 56, 1276–1285. 10.1016/j.neuroimage.2011.02.042

Logothetis, N.K., 2008. What we can do and what we cannot do with fMRI. Nature 453, 869–878. 10.1038/nature06976

Mareyam, A., Kirsch, J.E., Chang, Y., Madan, G., Wald, L.L., 2019. A 64-Channel 7T array coil for accelerated brain MRI, in: International Society for Magnetic Resonance in Medicine. p. 0764.

Markuerkiaga, I., Marques, J.P., Bains, L.J., Norris, D.G., 2021. An in-vivo study of BOLD laminar responses as a function of echo time and static magnetic field strength. Sci. Rep. 11, 1–13. 10.1038/s41598-021-81249-w

Norris, D.G., Polimeni, J.R., 2019. Laminar (f)MRI: A short history and future prospects. Neuroimage 197, 643–649. 10.1016/j.neuroimage.2019.04.082

Olman, C.A., Inati, S., Heeger, D.J., 2007. The effect of large veins on spatial localization with GE BOLD at 3 T: Displacement, not blurring. Neuroimage 34, 1126–1135. 10.1016/j.neuroimage.2006.08.045

Petridou, N., Siero, J.C.W., 2019a. Laminar fMRI: What can the time domain tell us? Neuroimage 197, 761–771. 10.1016/j.neuroimage.2017.07.040

Petridou, N., Siero, J.C.W., 2019b. Laminar fMRI: What can the time domain tell us? Neuroimage 197, 761–771. 10.1016/j.neuroimage.2017.07.040

Polimeni, Jonathan R., Fischl, B., Greve, D.N., Wald, L.L., 2010. Laminar analysis of 7T BOLD using an imposed spatial activation pattern in human V1. Neuroimage 52, 1334–1346. 10.1016/j.neuroimage.2010.05.005

Polimeni, Jonathan R, Fischl, B.R., Greve, D., Wald, L.L., 2010. Laminar-specific functional connectivity: distinguishing directionality in cortical networks, in: 16th Annu Meet Organ Hum Brain Mapp. p. 1402.

Polimeni, J.R., Renvall, V., Zaretskaya, N., Fischl, B., 2018. Analysis strategies for high-resolution UHF-fMRI data. Neuroimage 168, 296–320. 10.1016/j.neuroimage.2017.04.053

Polimeni, J.R., Wald, L.L., 2018. Magnetic Resonance Imaging technology — bridging the gap between noninvasive human imaging and optical microscopy. Curr. Opin. Neurobiol. 50, 250–260. 10.1016/j.conb.2018.04.026

Sirotin, Y.B., Hillman, E.M.C., Bordier, C., Das, A., 2009. Spatiotemporal precision and hemodynamic mechanism of optical point spreads in alert primates. Proc. Natl. Acad. Sci. U. S. A. 106, 18390–18395. 10.1073/pnas.0905509106

Tian, P., Teng, I.C., May, L.D., Kurz, R., Lu, K., Scadeng, M., Hillman, E.M.C., De Crespigny, A.J., D’Arceuil, H.E., Mandeville, J.B., Marota, J.J.A., Rosen, B.R., Liu, T.T., Boas, D.A., Buxton, R.B., Dale, A.M., Devor, A., 2010. Cortical depth-specific microvascular dilation underlies laminar differences in blood oxygenation level-dependent functional MRI signal. Proc. Natl. Acad. Sci. U. S. A. 107, 15246–15251. 10.1073/pnas.1006735107

van der Kouwe, A.J.W., Benner, T., Salat, D.H., Fischl, B., 2008. Brain morphometry with multiecho MPRAGE. Neuroimage 40, 559–569. 10.1016/j.neuroimage.2007.12.025

Wang, J., Nasr, S., Roe, A.W., Polimeni, J.R., 2022. Critical factors in achieving fine-scale functional MRI: Removing sources of inadvertent spatial smoothing. Hum. Brain Mapp. 43, 3311–3331. 10.1002/hbm.25867

Yu, X., Glen, D., Wang, S., Dodd, S., Hirano, Y., Saad, Z., Reynolds, R., Silva, A.C., Koretsky, A.P., 2012. Direct imaging of macrovascular and microvascular contributions to BOLD fMRI in layers IV-V of the rat whisker-barrel cortex. Neuroimage 59, 1451–1460. 10.1016/j.neuroimage.2011.08.001

Zaretskaya, N., Fischl, B., Reuter, M., Renvall, V., Polimeni, J.R., 2018. Advantages of cortical surface reconstruction using submillimeter 7 T MEMPRAGE. Neuroimage 165, 11–26. 10.1016/j.neuroimage.2017.09.060

